# A novel RNP compartment boosts translation in growing mouse oocytes to avoid cytoplasm dilution

**DOI:** 10.1101/2025.03.04.641373

**Authors:** N. Zollo, G. Zaffagnini, A. Canette, G. Letort, C. Da Silva, N. Tessandier, J. Dumont, C. Blugeon, S. Lemoine, B. Wattellier, E. Böke, M. Almonacid, M.-H. Verlhac

**Affiliations:** Center for Interdisciplinary Research in Biology (CIRB), Collège de France, Université PSL, CNRS, INSERM, 75005 Paris, France; Cologne Excellence Cluster for Aging and Aging-related Diseases (CECAD), University of Cologne, 50931 Cologne, Germany; Sorbonne Université, CNRS, Institut de Biologie Paris-Seine (IBPS), 75005 Paris, France; Department of Developmental and Stem Cell Biology, Institut Pasteur, CNRS UMR 3738, Université Paris Cité, 25 rue du Dr. Roux, 75015 Paris, France; ExposUM Institute, MIVEGEC, Université de Montpellier, IRD, CNRS, Montpellier, France; GenomiqueENS, Institut de Biologie de l’ENS (IBENS), Département de biologie, École normale supérieure, CNRS, INSERM, Université PSL, 75005 Paris, France; Phasics, Espace Technologique, 91190 Saint Aubin, France; Centre for Genomic Regulation (CRG), The Barcelona Institute of Science and Technology Barcelona, Spain; Institució Catalana de Recerca i Estudis Avançats (ICREA), Barcelona, Spain

## Abstract

Mammalian oocytes undergo a long growth phase in the ovary, during which transcriptional levels gradually decrease. Growing oocytes must therefore accumulate maternal stores and regulate their translation to achieve successful divisions and early embryo development. Using immunofluorescence, mass spectrometry and electron microscopy, we identified a novel and transient compartment, the Zollo Body, in late growing mouse oocytes, constituted of RNPs and organelles. Morphologically, this structure resembles the Balbiani body found in most vertebrate species but it stains positively for nascent translation and active phospho-mTOR. RNAseq analysis and dry mass measurements of growing oocytes with or without this compartment further support its key role in boosting translation, allowing growing oocytes to avoid cytoplasmic dilution despite their rapid size increase, ultimately ensuring their developmental potential.

## Introduction

Female fertility is at threat in modern societies and the cause can often be linked to poor oocyte quality. Oocytes are exceptionally big cells, and their quality and developmental potential are largely determined by the successful accumulation of maternal proteins, RNAs, lipids, and metabolites during their growth in the ovary. In mice, oocytes rely almost entirely on translational regulation during the latest stages of growth, when their size increase is suddenly boosted in a context of decreased transcription (*1–3*). The oocyte’s developmental potential therefore hinges on its ability to accumulate maternal mRNAs into ribonucleoproteins (RNPs) and fine-tune their translation.

In most vertebrates, including humans, a well-characterized structure called the Balbiani body is transiently assembled during the earliest stages of oogenesis before the growth phase, in primordial oocytes. The Balbiani body contains RNAs, RNPs but also ER, Golgi and mitochondria, and it is known to preserve these organelles and transcripts during early oogenesis (*4*, *5*). Proteomic profiling of the Balbiani body in *Xenopus* and zebrafish revealed a list of common proteins, including highly conserved ones involved in sequestering translationally silent mRNAs (*6*, *7*). The assembly of the Balbiani body in these two species is driven by a prion-like protein (XVelo/Bucky Ball homologs, respectively) capable of forming an amyloid-like matrix that reversibly traps organelles and RNPs (*6*, *8*). Both in *Xenopus* and zebrafish, XVelo/Bucky Ball proteins direct the incorporation of germline mRNAs into the Balbiani body, driving their insolubilization and consequently reducing their susceptibility to degradation (*9*). Fully grown mouse oocytes have also been found to form a membraneless condensate (mitochondrial-associated ribonuclear domain or MARDO), assembled via the RNP protein Zar-1 to silence maternal mRNAs (*10*). However, the presence of a proper Balbiani body in mouse oocytes has been recently challenged, as the structure identified in early mouse oogenesis does not contain mitochondria, a widely accepted hallmark of this compartment (*11*, *12*). So far, no other higher-level condensate similar to the Balbiani body from other species has been observed in mouse oocytes.

We describe a novel compartment, transiently assembled in late growing mouse oocytes, never characterized before. Its morphology reminds of the Balbiani body, as it is highly enriched in RNAs, RNPs, mitochondria and ER; this novel structure, however, appears much later in oogenesis, yet before the MARDO appearance (*10*). We used immunofluorescence, mass spectrometry and electron microscopy to systematically uncover its components and general organization. RNAseq analysis and dry mass measurements of growing oocytes with or without this compartment gave us insights into its role; interestingly, despite the similar appearance, this novel compartment seems to diverge from the canonical Balbiani body in terms of function, as it was found to be rather involved in translation. We confirmed active translation and enrichment in phospho-mTOR within the structure. Our results argue for a key role of this compartment in boosting translation to cope with the rapid size increase that mouse oocytes undergo towards the end of their growth, preventing cytoplasmic dilution known to be deleterious for cell fitness (*13, 14*).

## Results

### A novel compartment comparable to the Balbiani body

During the latest stages of their growth, mouse oocytes are enclosed into follicles that start forming a fluid-filled cavity called *antrum*. Oocytes retrieved from antral follicles were sorted according to their size and chromatin state as in (*3*) and to their zona pellucida thickness (Fig. 1A, Fig S1A and B). We identified a novel agglomerate of mitochondria among the late growing oocytes presenting an average diameter of 65 μm, at what we named the Cluster stage, before the Intermediary and Fully grown stages ((*3*); Fig. 1A and B). This structure forms alongside the nucleus with a variable size, its surface ranging from 50 to 800 μm^2^, corresponding to an average diameter of 21 μm (Fig. 1B). Importantly, this novel structure was not observed at any other stage of oocyte end of growth, suggesting that it is a transient structure disappearing with oocyte’s growth (Fig. 1A). This transient cluster is present in a subpopulation of late growing oocytes whose proportion varies as the animal ages (Fig. S1C). An accumulation of mitochondria close to the nucleus has been recently observed by another group, though it has not been further characterized (*15*). Other membranous organelles such as Golgi (Fig. 1C) and ER (Fig. 1D) are enriched in this compartment, reminding of the Balbiani body from other species (*16*). Furthermore, the cluster is enriched in RNAs (Fig. 1E) and stains positively for Thioflavin-T (Fig. 1F) indicating the presence of amyloid fibers, hallmark of proteins that contain a prion-like domain (PrLD) (*6*). Like for the zebrafish Balbiani body (*17*), microtubules seem to contribute to the cluster integrity, as treatment with Nocodazole led to its partial dissolution (Fig. 1G, left panel and Fig. S1E-F). On the contrary, F-actin is not involved in its maintenance since treatment with Cytochalasin D did not induce its disassembly (Fig. 1G, middle panel); microfilaments are not involved in its formation either, as proven by the presence of this cluster in oocytes invalidated for the actin nucleator Formin-2 (*18*, *19*), which lack cytoplasmic F-actin (Fig. 1G, right panel). An acentriolar MTOC (Microtubules Organizing Center), positive for Plk4 and Pericentrin, resides at the center of the cluster (Fig. 1H and I) where microtubules are nucleated (Fig. 1I and J, Fig. S1D). Upon Nocodazole washout, the cluster reformed progressively around the acentriolar MTOC, normally present at this stage (*20*), albeit slowly over a few hours (Fig. S1E-H), suggesting a potential role for microtubules in the assembly of this compartment. Altogether, our observations disclose the presence of a novel transient structure, that we named the Zollo body (Zb), whose composition resembles the Balbiani body, assembled and maintained in part by microtubules.

**Fig. 1.**
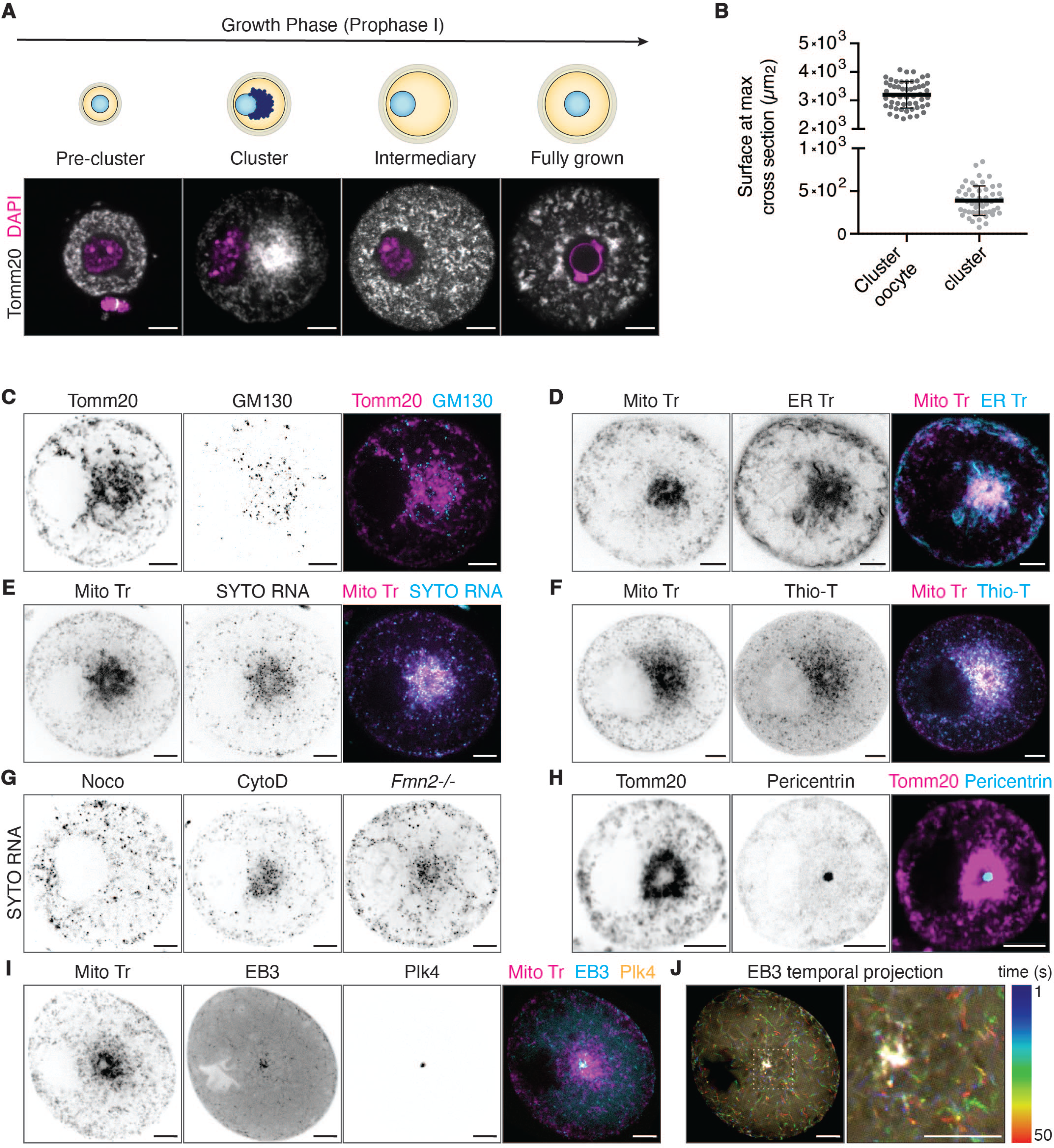
Antral growing mouse oocytes contain a novel cluster of organelles and RNPs. **(A)** Immunofluorescence staining of mouse oocytes displaying mitochondria labeling (Tomm20, white) at different stages of the end of the follicular growth phase, sorted according to their size and chromatin state (DAPI, pink) as in (*3*). **(B)** Quantification of oocyte and cluster surface, measured at their maximal cross section in Cluster stage oocytes. N=53 for both oocyte and cluster measurements; 3 separate experiments. **(C)** Immunofluorescent staining indicating enrichment of Golgi (GM130, middle) in the cluster, identified by labeling mitochondria (Tomm20, left). The right panel shows the merge of mitochondria (pink) and Golgi (blue) staining. **(D)** Cluster stage oocyte stained with Mito Tracker (Mito Tr, left) and ER Tracker (ER Tr, middle). The right panel shows the merge of Mito Tr (pink) and ER Tr (blue) staining. **(E)** Cluster stage oocyte stained with Mito Tr (left) and SYTO RNA Select (SYTO RNA, middle) to detect mRNAs localization. The right panel shows the merge of Mito Tr (pink) and SYTO RNA (blue) staining. **(F)** Cluster stage oocyte stained with Mito Tr (left) and Thioflavin-T (Thio-T, middle) to reveal the presence of beta-amyloid fibrils. The right panel shows the merge of Mito Tr (pink) and Thio-T (blue) staining. **(G)** Cluster stage oocyte treated with Nocodazole (Noco, left), Cytochalasin D (Cyto D, middle) or collected from *Fmn2^-/-^* mice (right), stained with SYTO RNA Select. **(H)** Cluster stage oocyte stained for mitochondria (Tomm20, left) and Pericentrin (middle) to visualize the MTOC. The right panel shows the merge of Tomm20 (pink) and Pericentin (blue) staining. **(I)** Cluster stage oocyte stained with Mito Tr (left) and co-expressing EB3- GFP (EB3, middle left) to visualize microtubules growing (+) ends and mcherry-Plk4 (Plk4, middle right) to visualize MTOC. The right panel shows the merge of Mito Tr (pink), EB3 (blue) and Plk4 (yellow) staining. **(J)** Temporal color-coded projection of a stream movie on EB3-GFP (left) and its zoom (dashed white square, right); acquisition every 500 ms for 50 s. Data are shown as mean ± SD. Scale bars are 10 μm.

### Proteomic profiling reveals enrichment in ribosomes

We were able to mechanically isolate the Zollo body, which remained stable for several hours outside of the cell (Fig. 2A and B), as previously observed for the Balbiani body in *Xenopus* and zebrafish (*6*, *7*), even allowing for fixation on glass coverslips and immunostaining (Fig. 2C). The isolated Zb was submitted to mass spectrometry, using the whole oocyte at the same Cluster stage as a control. This analysis identified 2167 proteins in the control whole oocyte and 585 in the isolated Zb, with 42 proteins being enriched in the isolated Zb compared to the control (Fig. 2D and Table 1). Gene ontology on the 585 proteins of the isolated Zb revealed a surprising predominance in ribosomal subunits, along with an expected high representation of proteins from membranous organelles and from mitochondria (Fig. 2E). A closer look at the 42 proteins whose relative abundance was higher in the isolated Zb (Fig. 2F) showed the presence of proteins involved in the translation machinery and of few proteins with PrLDs (underlined in Fig. 2F, Fig. S2A), consistent with the Thioflavin-T staining (Fig. 1F). Using immunofluorescence, we notably confirmed the enrichment of Ddx3x, a helicase with a PrLD, inside the Zb (Fig. 3A and C, Fig. S2A). Such PrLD-containing proteins could play a role in the assembly of this novel compartment, similarly to XVelo/Bucky ball in the Balbiani body of *Xenopus* and zebrafish respectively (*6*, *7*). Consistent with this idea, and with the fact that the Zb could be isolated mechanically, Ddx3x-GFP presented an incomplete and slow recovery of fluorescence after photobleaching inside the Zb, while it recovered fully and rapidly in the cytoplasm (Fig. 3D and E). This result suggests that Ddx3x could be part of a structural amyloid matrix participating in the cohesivity of the Zb.

**Fig. 2.**
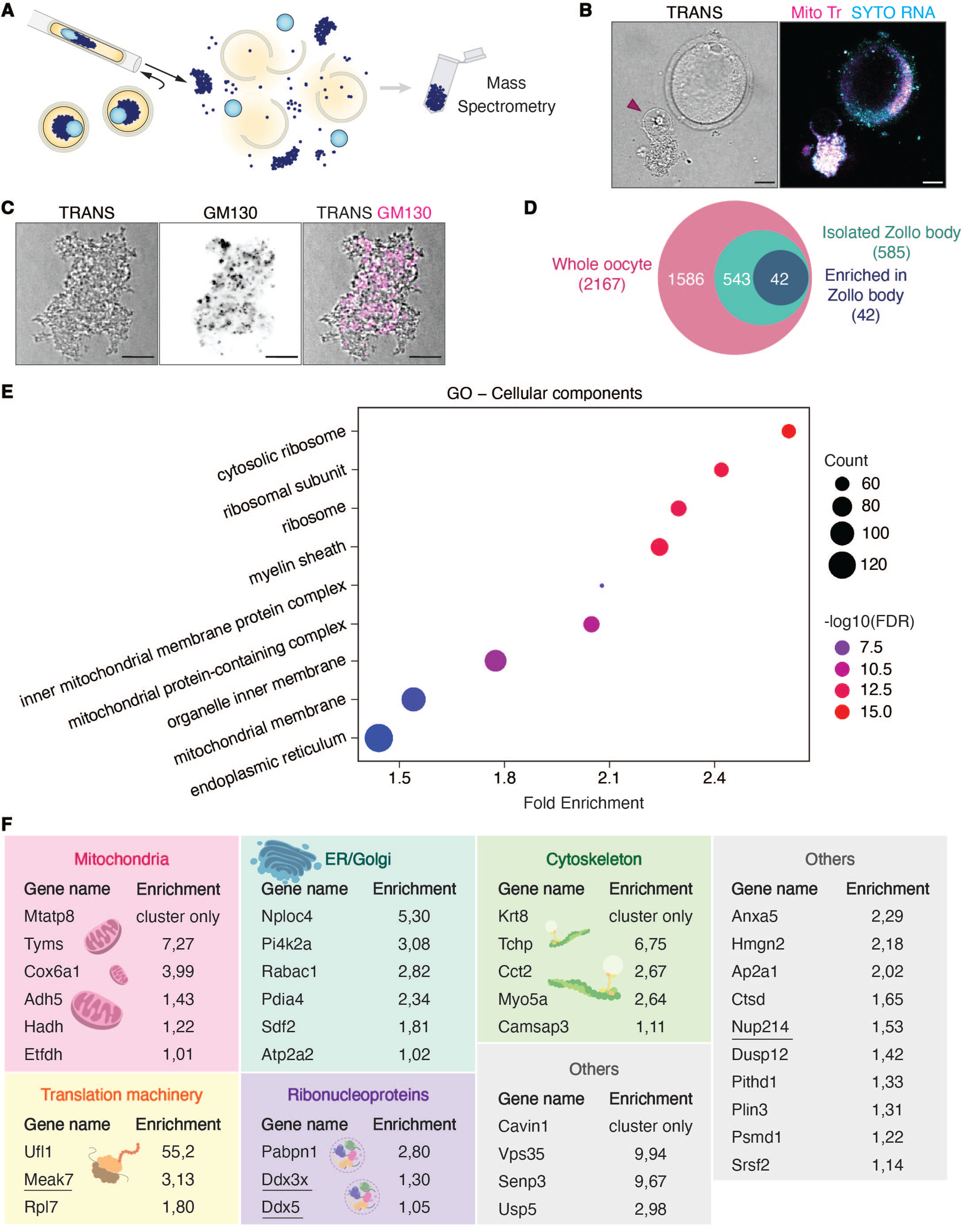
Mass spectrometry analysis of isolated Zollo bodies reveals an enrichment in ribosomes. **(A)** Schematic representation of the Zollo body’s mechanical isolation for mass spectrometry analysis. **(B)** Transmitted light image of a mechanically isolated Zb and its corresponding remaining oocyte (TRANS, left), co-stained with Mito Tr (pink) and SYTO RNA (blue, right panel); the purple arrowhead indicates the nucleus attached to the cluster. **(C)** Transmitted light image of an isolated Zb (TRANS, left), stained for GM130 (GM130, middle) to visualize Golgi complexes. The right panel shows the merge of TRANS (grey levels) and GM130 (pink). **(D)** Diagram showing relative proportions of proteins identified by mass spectrometry analysis in the whole oocyte at the Cluster stage (pink), in the isolated Zb (green) and enriched in the isolated Zb (blue). The numbers refer to the total number of proteins for each category (whole oocyte, isolated cluster and enriched in the cluster). **(E)** Enriched GO pathways for proteins identified in the isolated Zb. **(F)** Schematized list of the 42 proteins found enriched in the isolated Zb; each gene is shown with its corresponding enrichment value, obtained from the ratio of the normalized abundance in the isolated Zb over the abundance in the whole oocyte at the Cluster stage. Proteins with a prion-like domain are underscored. Scale bars are 10 μm.

**Fig. 3.**
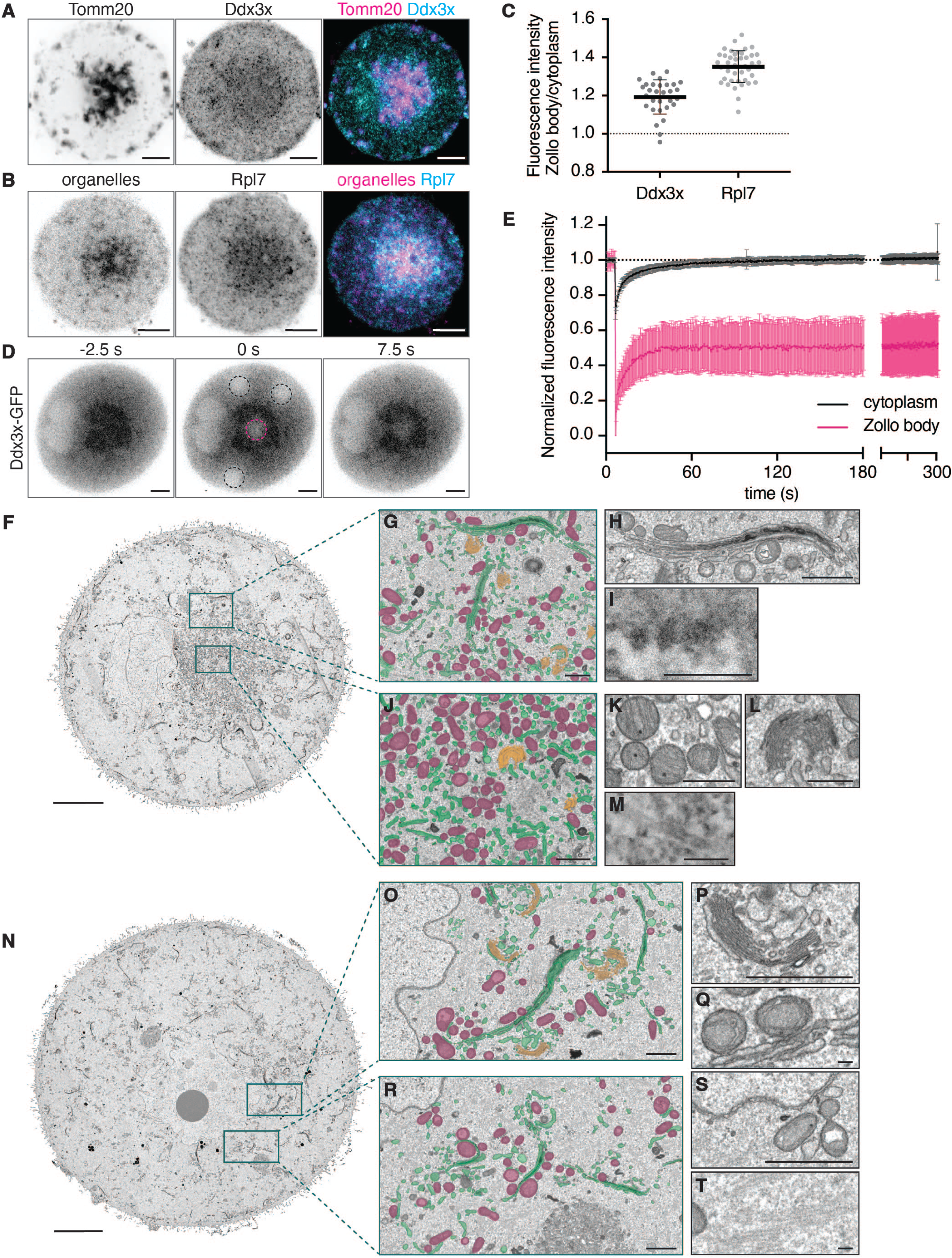
The Zb is packed with RNPs, ribosomes and organelles in a solid-like structure. **(A)** Cluster stage oocyte stained for Tomm20 (left) and Ddx3x (middle). The right panel shows the merge of Tomm20 (pink) and Ddx3x (blue) staining. **(B)** Cluster stage oocyte stained to visualize the ribosomal subunit Rpl7 (middle); the Zb is visualized via autofluorescence of its organelles at 488 nm (organelles, left). The right panel shows the merge of mitochondria (pink) and Rpl7 (blue) staining. **(C)** Quantification of Ddx3x and Rpl7 signal intensities in the cluster compared to the cytoplasm for (A) and (B). N= 30 for Ddx3x, 4 separate experiments; N= 42 for Rpl7, 3 separate experiments. **(D)** FRAP experiment performed on Cluster stage oocyte expressing Ddx3x-GFP. Dotted pink and black circles are regions of interest bleached respectively inside the Zb and inside the surrounding cytoplasm. **(E)** Quantification of the FRAP experiment in (D). The pink and black recovery curves correspond to regions in the Zb and in the cytoplasm, respectively. N= 21; 3 separate experiments. **(F)** Electron Microscopy image of a Cluster stage oocyte and of **(N)** an Intermediary stage oocyte; insets are magnifications of outlined regions. In (G), (J), (O), (R) color coding was applied to mitochondria (purple), Golgi ministacks (orange), ER (green). Insets (H), (I) are magnifications from inset (G); insets (K), (L), (M) are magnifications from inset (J); insets (P), (Q) are magnifications from inset (O); insets (S), (T) are magnifications from inset (R). Scale bars for (A), (B), (D), (F), (N) are 10 μm; scale bars for (G), (J), (O), (R), (P), (S) are 1 μm; scale bars for (H), (K), (L) are 500 nm; scale bars for (I), (M), (Q), (T) are 100 nm.

**Table 1.** Mass Spectrometry analysis on isolated Zb vs Cluster stage oocytes. About 50-60 isolated Zbs were processed (sample S1=100% of total submitted sample), with 50 whole oocytes at the Cluster stage as control (sample S2=10% of total submitted sample; sample S3=20% of total submitted sample). Second sheet: raw data as provided by the platform. Third sheet: result of the enrichment analysis as described in methods. Table 1 refers to Fig. 2.

Using immunofluorescence, we further showed the enrichment in the Zb of ribosomal proteins (Fig. 3B and C) as well as of Keratin 8 (Fig. 2F, Fig. S2B and D) highlighted by mass spectrometry. While the Zb contained RNPs such as Ddx3x, it was not enriched in the canonical P-body marker Lsm14a (Fig. S2C and D) (*21*, *22*).

Together, we find that the Zb is enriched in ribosomal and mitochondrial proteins and has material properties consistent with a solid-like compartment.

### Electron microscopy confirms overall Zb organization

To better characterize the overall organization inside the novel compartment identified in late growing oocytes, we performed electron microscopy on Cluster and Intermediary stage oocytes (see Methods). Consistent with live, immunofluorescent and proteomic data, the Zb was observed at the center of the oocyte, located next to the nucleus (Fig. 3F). The structure displayed a clear enrichment in ER (green in Fig. 3G, J and enlarged in Fig. 3H), in mini-Golgi stacks (orange in Fig. 3G, J and enlarged in Fig. 3L) and was extremely dense in mitochondria (purple in Fig. 3G, J and enlarged in Fig. 3K). Ribosomes could be spotted along the membrane of the ER (Fig. 3I). The center of the Zb showed a region less dense in organelles (Fig. 3J), as observed by live and immunofluorescent staining (Fig. 1D-I), where microtubules were observed (Fig. 3M). This novel compartment was no longer visible in Intermediary oocytes (Fig. 3N), as observed in Fig. 1A, and the cytoplasm presented a more dispersed distribution of mitochondria (Fig. 3O, R), ER (Fig. 3O, R, Q), mini-Golgi stacks (Fig. 3O, R, P), even in regions close to the nuclear envelope (Fig. 3S). Cytoplasmic lattices were also observed (Fig. 3T). The EM data validated the global architecture of the Zb, and further confirmed its enrichment in mitochondria and ribosomes, as identified using mass spectrometry and immunofluorescence on the isolated structure.

### RNA sequencing comparing Cluster and Intermediary oocytes points towards a function in translation

As the Zb contains RNA (Fig. 1E) and RNPs (Fig. 2F and 3A), we decided to perform RNA sequencing and compare Cluster with Intermediary oocytes to identify differentially expressed RNAs between the two stages. The sequencing identified 252 and 407 RNAs respectively significantly (p<0.05) down and up-regulated at the Cluster stage compared with the Intermediary stage (Fig. 4A, Table 2). Gene set enrichment analysis (GSEA) suggests an up-regulation in the Cluster stage of genes associated with GTPase activity, helicase activity as well as DNA-binding negative regulators of transcription (Fig. 4B). The enhanced GTPase activity, particularly for regulators of the actin cytoskeleton, is consistent with the important changes in the actin mesh organization occurring between the Cluster and Intermediary stage (*3*). Similarly, the DNA-binding negative regulators of transcription category is in line with the transcriptional silencing that follows the Cluster stage (*1*, *2*). The enrichment in helicase activity fits well with the presence of Ddx5 and Ddx3x as identified by the proteomic data on the isolated Zb (Fig. 2F and 3A). The analysis of the down-regulated genes at the Cluster stage highlights two major functions, translation and mitochondrial activities (Fig. 4C and 4D). Although it may seem counterintuitive that these genes are down-regulated at the Cluster stage while ribosomal and mitochondrial proteins are enriched in the Zb, this could indicate an intensive translation of mRNAs and maybe increased turn-over of these specific gene sets during the Cluster stage. In eukaryotes, mRNAs encoding ribosomal proteins present a half-life of 9h (*23*), compatible with the estimated duration of a few days between the Cluster and Intermediary stages.

**Fig. 4.**
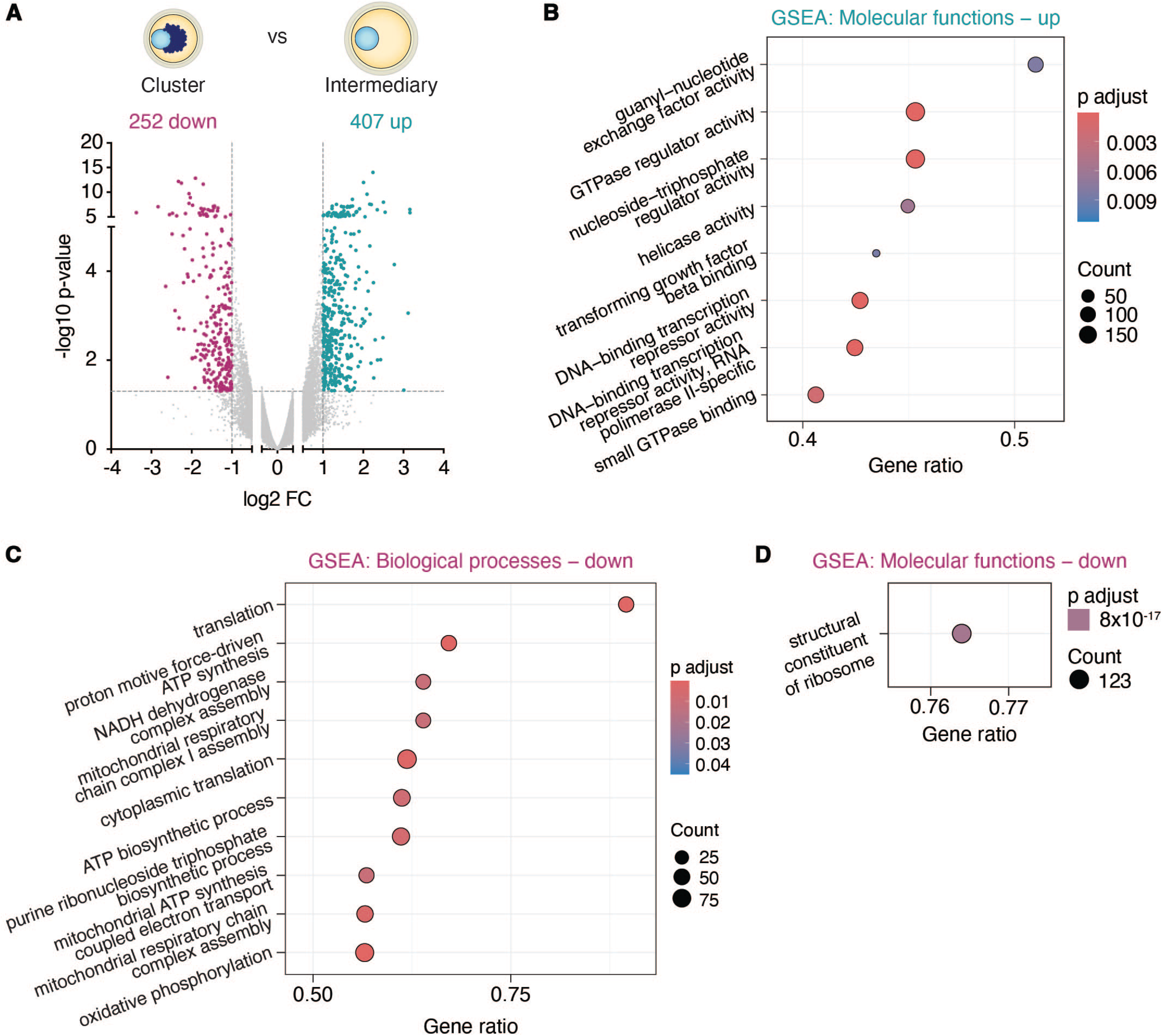
RNAseq analysis suggests a role in translation. **(A)** Volcano plot depicting differentially expressed genes at Cluster versus Intermediary stage oocytes. 252 down and 407 upregulated genes (purple and green dots respectively), with an adjusted p- value<0.05 and average log2 fold change of <-1 and >1. **(B)** Gene Set Enrichment Analysis on upregulated genes from (A) showing enriched GO pathways in the category Molecular functions. **(C)** Gene Set Enrichment Analysis on downregulated genes from (A) showing enriched GO pathways in the category Biological processes and **(D)** Molecular Functions.

**Table 2.** RNA sequencing results: Cluster vs Intermediary stage oocytes. Second sheet: raw data obtained from DEseq analysis. Third sheet (pink): list of downregulated RNAs with a log2 FC < -1 and p value < 0,05. Fourth sheet (green): list of upregulated RNAs with a log2 FC > 1 and p value < 0,05. Table 2 refers to Fig. 4.

### Multiple evidence arguing for active translation inside the Zb

To test whether the Cluster stage is associated with active translation, we first used a Click-iT chemistry technology that employs a modified fluorescent puromycin probe to detect nascent peptides ((*24*) and methods). We observed a significant accumulation of fluorescent labeling inside the Zb (Fig. 5A) that was abrogated by addition of the non- fluorescent competitor puromycin, arguing for its specificity (Fig. 5A, quantified in 5C). This observation provided direct evidence of active translation inside the novel compartment. Activation of the mTOR pathway, and in particular phosphorylation of mTORC1, has been shown to integrate various metabolic signals and promote pathways such as ribosome biogenesis and protein synthesis, eventually leading to cell growth (*25*). As second evidence for active translation, we detected an enrichment of phosphorylated- mTOR inside the Zb compared to the surrounding cytoplasm (Fig. 5B, quantified in 5D). The overall levels of activated mTORC1 decreased upon treatment with Nocodazole (Fig. 5E, quantified in 5F), highlighting the importance of Zb integrity to maintain translational activity. As last evidence, we measured the dry mass at the Cluster, Intermediary and Fully grown stages of oocyte growth. Results highlighted a substantial increase in oocyte diameter between the Cluster and Intermediary stage while their growth was not significant later on (Fig. 5G). Using quantitative phase imaging (methods) we observed a corresponding substantial increase in dry mass between the Cluster and Intermediary stages (Fig. 5H). Dry mass is a good proxy to estimate the levels of protein synthesis, since proteins constitute 60% of the dry mass of mammalian cells (*26*). These results strongly argue for sustained protein synthesis between the Cluster and Intermediary stages. Importantly, based on previous literature (*3*, *27–31*), we could establish that oocytes at this stage experience an acceleration in volume increase (Fig. S3A and B); a massive accumulation of proteins is therefore fundamental to prevent the decrease in oocyte cytoplasmic density that would otherwise be associated to such a fast growth (Fig. 5I). While dry mass increases linearly with oocyte diameter for the three stages (Fig. 5J), quantitative phase imaging allowed to reveal that the Cluster stage is peculiar when it comes to protein density. Compared to later stages, the Cluster stage preserves a constant cytoplasmic density no matter the size of the oocyte, denoting a tighter control over protein content (Fig. 5K). Altogether, these observations advocate for active translation inside the Zb, boosting oocyte growth at a crucial stage before the shut-down of transcription.

**Fig. 5.**
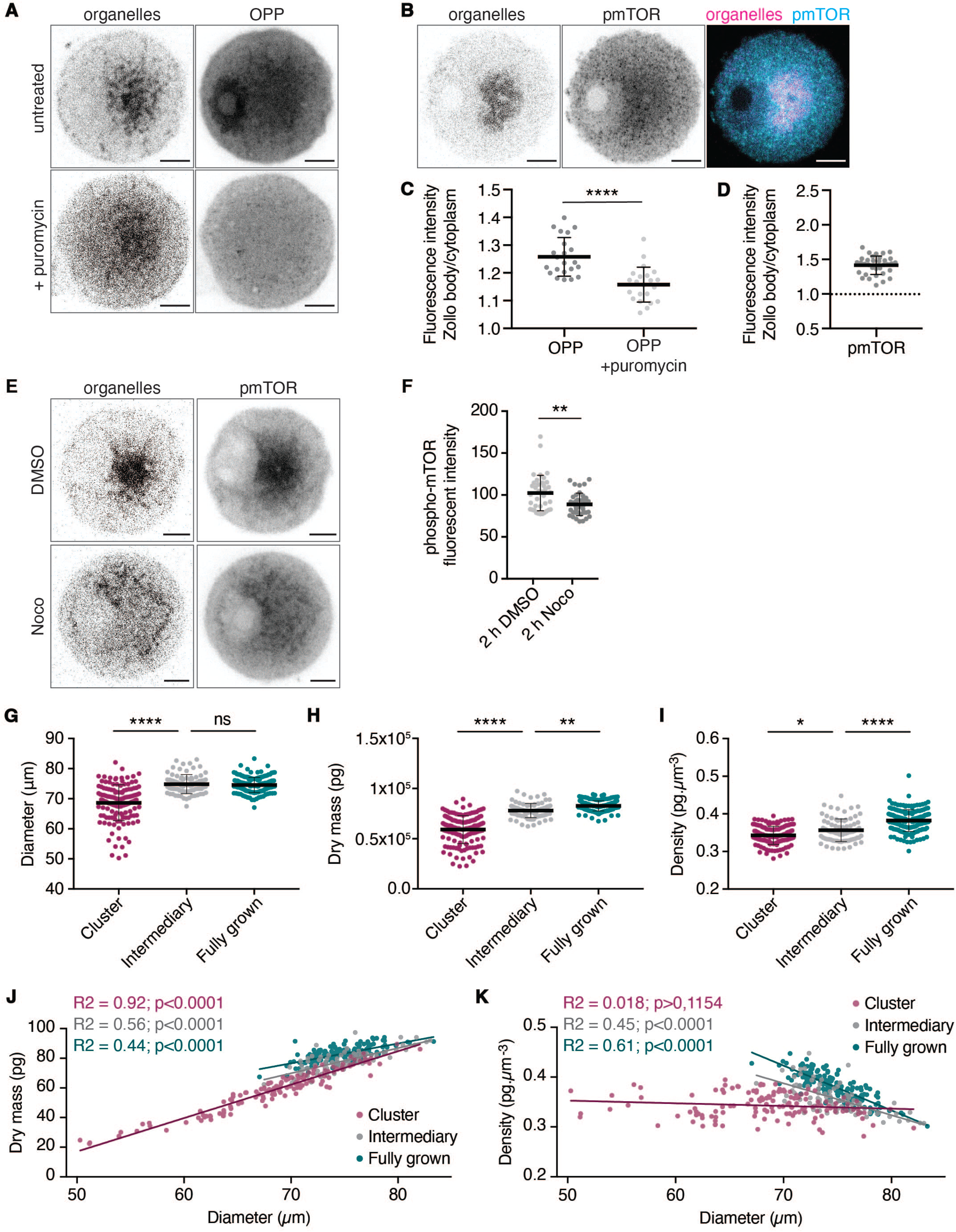
The Zb is associated with active translation. **(A)** Oocyte at the Cluster stage stained for nascent translation by Click chemistry with the puromycin analog OPP (OPP, right column), negative control condition previously treated with puromycin (+ puromycin, bottom row). The Zb is visualized through the autofluorescence of its organelles at 488 nm (organelles, left column). **(B)** Cluster stage oocyte labelled with anti-Phospho-mTOR Ser2448 (pmTOR, middle); the Zb is visualized through the autofluorescence of its organelles at 488 nm (organelles, left). The right panel shows the merge of organelles (pink) and pmTOR (blue) staining. **(C)** Quantification of nascent translation signal intensity at Cluster stage as in (A). N= 22 for OPP, N= 22 for OPP+puromycin, 5 separate experiments. **(D)** Quantification of Phospho-mTOR signal intensity at Cluster stage as in (B). N= 34, 4 separate experiments. (**E)** Oocyte at the Cluster stage labelled with anti- Phospho-mTOR Ser2448, (right column) after 2hrs treatment with Nocodazole (Noco, bottom row) or its solvent DMSO (top row). The Zb is visualized by autofluorescence of its organelles at 488 nm (organelles, left column). **(F)** Quantification of fluorescence intensity at Cluster stage from (E). N= 39 for DMSO 2 h, N= 39 for Nocodazole 2 h, 5 separate experiments. **(G)** Oocyte diameter at Cluster, Intermediary and Fully grown stages measured with quantitative phase imaging. N= 139 for Cluster, N= 82 for Intermediary, N= 160 for Fully grown; 6 separate experiments. **(H)** Oocyte dry mass at Cluster, Intermediary and Fully grown stages measured with quantitative phase imaging. N= 139 for Cluster stage, N= 82 for Intermediary stage, N= 160 for Fully grown stage; 6 separate experiments. **(I)** Oocyte protein density at Cluster, Intermediary and Fully grown stages estimated from measurements in (G) and (H). N= 139 for Cluster, N= 82 for Intermediary, N= 160 for Fully grown; 6 separate experiments. **(J)** Simple linear regression of dry mass values from (H) plotted against diameter measurements from (G) for Cluster, Intermediary and Fully grown stages. **(K)** Simple linear regression of density values from (I) plotted against diameter measurements from (G) for Cluster, Intermediary and Fully grown stages. Data are shown as mean ± SD. P values were calculated using unpaired two-tailed Student’s t test with Welch’s correction for (C), Mann-Whitney test for (F) or one-way ANOVA with Tukey’s post hoc test for (G), (H), (I). Scale bars 10 μm. ns = not significant; * = p<0.05; ** = p<0.01; *** = p<0.001; **** = p<0.0001.

## Discussion

The global architecture of the novel compartment here characterized, that we named Zollo body, is reminiscent of the Balbiani body. In fact, it contains RNAs, RNPs, mitochondria, Golgi, ER and is organized around a MTOC that nucleates microtubules despite being acentriolar. It is also most probably held together by an amyloid-matrix, inside which diffusion appears slowed down. However, the Zollo body is well distinguished from the Balbiani body. First, the Zollo body appears in late oogenesis while the Balbiani body is formed early on, in primordial oocytes (*16*). Second, the Balbiani body has been associated with quiescence and storage of maternal mRNAs as well as preservation of organelles, while we provide multiple evidence that the Zollo body is associated with active translation and appears at a stage where growth is accelerated. Increasing evidence in the germline of several organisms supports a role in localized translation for RNPs condensates (*32*); the Zollo body would therefore constitute a novel example of higher-level condensate promoting translation. The Zollo body’s function in promoting translation is also in line with the presence of the Ddx3x helicase as part of the amyloid matrix structuring the cluster, since Ddx3x is involved in the translation of mRNAs with complex 5’UTRs (*33*). This function could be of key importance to the end of oogenesis when transcription is progressively switched off (*1–3*). In this context, we propose that this structure could avoid dilution of the cytoplasm while the oocyte undergoes a massive increase in size, preventing the oocyte from experiencing senescence, as observed in other cell types (*13*, *14*). This could be relevant considering that Fully grown oocytes closer to their meiotic divisions appear less efficient in adjusting their density with respect to their size. Therefore, the Zollo body strategically emerges at a critical stage of oogenesis, just before the highly asymmetric meiotic divisions that help preserving the maternal stores in the cytoplasm (*34*).

It is striking for a solid-like structure such as the Zollo body to be associated with an active process like translation. Solid amyloid structures are hallmarks of pathologies such as neurodegenerative diseases, and the formation of ectopic solid-like condensates impairs oogenesis in *C. elegans* (*35*). On the other hand, the Balbiani body is associated with dormancy (*36*). One hypothesis is that the Zollo body might be porous, containing micro- compartments that allow free molecular diffusion. Confinement within such compartments could even enhance enzymatic reactions, with pores increasing the surface area available for these processes. In this context, mitochondria trapped within the matrix may not just be protected but could also serve as a source of ATP for enzymatic reactions, especially considering that protein synthesis is the most ATP-consuming process in growing cells (*37*). Notably, Ddx3x, a protein involved in translation, functions as an ATP-dependent helicase.

What maintains the cohesion of the Zb remains an open question. Since rodents homologs of Buc from zebrafish Balbiani body (*38*) and XVelo from *Xenopus* (*6*) have not been identified so far (*38*), it seems rather unlikely that a sole “glue” protein contributes to the integrity of the cluster. Instead, several prion-like proteins within the cluster, such as Ddx3x, may be involved, along with microtubules and Keratin 8. Interestingly, the *Xenopus* Balbiani body is similarly enriched in Keratin 18, albeit to a lesser extent (*6*).

Ddx3x condensates have been reported in several physiological contexts, for example as part of stress granules (*39*) or playing a key role during the maternal to zygotic transition in zebrafish (*40*). In parallel, several cancers and neurodegenerative disorders have been correlated to Ddx3x mutants (*41*, *42*). However, apart from one Ddx3x pathological mutant that forms amyloid-like fibrils (*42*), other described Ddx3x condensates display liquid properties. Here, the slow diffusivity of Ddx3x within the Zollo body suggests that Ddx3x contributes to its amyloid-like properties, though it remains possible that Ddx3x transitions from a liquid to a solid state, consistent with the hypothesis of a porous matrix.

It is remarkable how the Zollo body in mouse oocytes, a Mitochondria-associated RNP condensate like the MARDO (*10*), yet structurally more reminiscent of the Balbiani body from other vertebrates, serves a previously unreported function for these type of amyloid- like aggregates, and at a different stage of oogenesis. This is particularly interesting as we noticed from published datasets that both the zebrafish and *Xenopus* Balbiani body also contain ribosomes and other translation-related factors (*6*, *7*). From an evolutionary point of view, the comparison between the Zollo body and the Balbiani body of other species is particularly fascinating, as it suggests that a similar structure, composed of the same components, can perform vastly different functions depending on the context.

By proposing an unreported role for a Balbiani body-like compartment in mouse oocytes, this work broadens the functional repertoire of this type of structure and, as suggested in (*5*), could reveal their versatility in female germ cells.

## Acknowledgments

We thank ME Terret for critical reading of the manuscript and all members of the Verlhac-Terret laboratory for discussions, the CIRB animal and Orion facilities. The Verlhac-Terret laboratory is supported by CNRS, INSERM, Collège de France and the Bettencourt Schueller Foundation. This work received support under the program « Investissements d’Avenir » launched by the French Government and implemented by the ANR, with the references: ANR-10-LABX-54 MEMO LIFE, ANR-11-IDEX-0001-02 PSL Research University. NZ was supported by fellowships from French Ministère de la Recherche and from ARC (ARCDOC42024010007657). E.B. and G.Z acknowledge funding from the Ministerio de Ciencia e Innovacio no PID2020-115127GBI00. This work was supported by the France Génomique national infrastructure, funded as part of the “Investissements d’Avenir” program managed by the Agence Nationale de la Recherche (contract ANR-10-INBS-0009).

## Author contributions

Conceptualization: MHV, MA

Methodology: MHV, MA, EB, NZ

Investigation: NZ, GZ, AC, GL, CDS, NT, JD, CB, SL, BW

Visualization: NZ, GZ, AC, GL, CDS, NT, CB, SL

Funding acquisition: MHV, MA

Project administration: MHV

Supervision: MHV, MA

Writing – original draft: MHV, NZ, MA

Writing – review & editing: MHV, NZ, MA, GZ, EB, AC

## Materials and Methods

### Mouse oocyte collection

All animal studies were performed in accordance with the guidelines of the European Community and were approved by the French Ministry of Agriculture (authorization N°75– 1170) and by the Direction Générale de la Recherche et de l’Innovation (DGRI; GMO agreement number DUO-5291). Mice were housed in the animal facility on a 12-h light/dark cycle, with an ambient temperature of 22–24°C and humidity of 40–50% and received food and water *ad libitum*. Mice used in this study include female C57BL/6J (Charles River Laboratories; 3 to 17 weeks old), Formin 2 knockout female mice (*Fmn2 -/-*) (*43*) (8 weeks old). At least three mice were used per experiment. Ovaries were extracted from mice as previously described (*44*) into pre-warmed at 37°C M2 + Bovine Serum Albumin (BSA; Sigma, A3311) medium supplemented with 1 μM Milrinone (Merck, M4659) as in (*45*), which maintains growing oocytes arrested in Prophase I. Ovarian follicles were punctured with surgical needles to release growing oocytes from antral follicles (end of oocyte growth). All subsequent oocyte culture and live imaging steps were then carried out in M2+BSA+1 μM Milrinone under oil at 37°C (Sigma, M8410).

### Immunofluorescence

Oocytes were treated with 0.4% Pronase (Sigma, P5147) to remove the zona pellucida. Then they were fixed in 4 % paraformaldehyde at 30 °C for 30 min on coverslips coated with gelatin and poly-L-lysine (*3*). Oocytes were permeabilized and pre-blocked for 15min in PBS with 0.5% Triton-X and 3% BSA. Oocytes were extensively washed with PBS buffer between solutions. Oocytes were incubated with primary antibodies overnight at 4 °C and with secondary antibodies for 1 hour at room temperature. Coverslips were then mounted on glass microscope slides in 250nm thick chambers (Electron Microscopy Sciences, 70366-12) to avoid oocyte squashing, using ProLong Gold antifade reagent with DAPI (Thermofisher, P36931). Primary and secondary antibodies were diluted in PBS with 0.2% Triton-X and 3% BSA. Incubation with anti-Phospho-mTOR was in PBS with 3% BSA only. Isolated Zollo bodies were not permeabilized and incubations with antibodies were in PBS with 0.05% Triton X-100 and 3% BSA. The following antibodies were used: rabbit anti-Tomm20 (1:800; Abcam, ab186735), mouse anti-Tomm20 (1:1200; Novus, H9804-M01), mouse anti-GM130 (1:100; BD, 610822), mouse anti-Pericentrin (1:400; BD, 611814), mouse anti-Ddx3 (1:800; Proteintech, 67915), rabbit anti-Rpl7 (1:400; Sigma Atlas, HPA046794), rabbit anti-Phospho (Ser2448)-mTOR (1:800; Cell Signalling, 5536S), rat anti-TROMA1 (Keratin 8, 1:800; DHSB, AB531826), rabbit anti-Lsm14a (1:500; Thermofisher, PA5-78464) and species-specific Alexa Fluor secondary antibodies (1:400, Thermo Fisher).

### Live markers and drugs

MitoTracker Deep Red (ThermoFisher, M22426) was diluted in DMSO to make a 1 mM stock solution, used at 100 nM for 30 minutes; ER Tracker Red (ThermoFisher, E34250) was diluted in DMSO at 1 mM, used at 1 µM for 20 minutes; SiR-Tubulin (Spyrochrome, SC002) was diluted in DMSO to make 1 mM stock solution, used at 5 µM for 20 minutes. SYTO RNA Select (ThermoFisher, S32703) was diluted in DMSO to make a 5 mM stock solution, used at 5 µM for 15 minutes; Thioflavin-T (Abcam, ab120751) was diluted in DMSO to make a 100 mM stock solution, used at 10 µM for 15 minutes. Puromycin (Sigma, P7255) was diluted in water to make a 50 mg/mL stock solution, used at 100 µg/ml for 1 hour. Cytochalasin D (Sigma, C8273) was diluted in DMSO at 10 mg/ml, used at 1 µg/ml for 30 minutes. Nocodazole (Sigma, M1404) was diluted in DMSO at 10 mM, used on oocytes at 1 µM for 30 minutes. Culture medium containing live markers or drugs at their working concentration was freshly prepared and prewarmed at 37°C. All stock solutions and aliquots were stored at −20°C.

### Microinjection

Oocytes were microinjected with cRNAs using an Eppendorf Femtojet microinjector. Oocytes were kept in Prophase I between 2 and 4 hours to allow expression of fusion proteins. All live culture and imaging were carried out under oil at 37°C. We used the following constructs: mcherry-Plk4 (*46*), EB3-GFP (*47*) and Ddx3x-GFP (this study). pRN3-Ddx3x-GFP was obtained by subcloning Ddx3x ORF clone (CliniSciences, MG226484) into the EcoRI-Sall sites of a pRN3-GFP vector (Genscript). *In vitro* synthesis of capped cRNAs was performed as previously described (*48*). cRNAs were centrifuged at 4 ◦C for 45 min at 13,000 rpm before microinjection.

### Live imaging

Spinning-disk images were acquired at 37°C using a Plan-APO ×40/1.25 NA objective on a Leica DMI6000B microscope enclosed in a thermostatic chamber (Life Imaging Service) equipped with a Retiga 3 CCD camera (QImaging, Burnaby) coupled to a Sutter filter wheel (Roper Scientific) and a Yokogawa CSU-X1-M1 spinning-disk. Metamorph software (Molecular Devices) was used to collect data. To follow EB3-GFP labelling of microtubules, oocytes were imaged every 500 ms with the stream acquisition mode of Metamorph upon 491 nm excitation. All other live and fixed samples were acquired with 3 µm z-step over a slice 27 µm thick. Time-lapse images of oocytes after Nocodazole washout were acquired every 20 or 30 min. Exposition was 500 ms in all channels for all acquisitions.

### Mass Spectrometry and enrichment analysis

A total of 14 C57BL/6J female mice (Charles River laboratories) aged four weeks were used for the experiments. Prophase I arrested oocytes were collected in M2+ PVA medium supplemented with 1 μM Milrinone. Clean glass pipettes with a diameter of about 20 µm were used to disrupt Cluster stage oocytes and isolate the novel structure. Between 50 and 60 isolated Zollo bodies or 50 whole oocytes at the Cluster stage were collected in 1 µl medium, transferred into 15µl of 6M Guanidinium Chloride and stored at -80°C until processing. Samples were processed for mass spectrometry and analyzed as described (*49*). Results in Table 1: sample S1 contained all collected isolated Zollo bodies, while samples S2 and S3 contained respectively 10% and 20% of the whole Cluster stage oocyte sample. Raw data was filtered to remove contaminants (“CON” in Accession column) and to keep only proteins identified with a high confidence (Protein FDR Confidence column). Since the total protein content was different in each sample, the Relative Abundance of each protein was calculated as a percentage in the three samples: (Abundance value of a protein in a sample S / total protein content of S)*100, where the total protein content corresponds to the sum of Abundance values of all proteins found in the sample. The ratio between the Relative Abundance from Sample 1 and 3 was then calculated. To identify the proteins enriched in the isolated Zb, a filter was applied to the “Found in Sample S1” column to remove the “Not Found” occurrence; proteins were then ordered in decreasing value of ratio between Relative Abundances of Sample 1 and Sample 3. Proteins found in the isolated Zb whose ratio in Relative Abundance compared to Sample 3 was >1 were considered enriched in the isolated Zb. To check for PrLDs, protein sequences were obtained from UniProt (https://www.uniprot.org/id-mapping) and uploaded on PLAAC website (Prion-like Amino Acid Composition, (*50*) http://plaac.wi.mit.edu/).

### Fluorescence recovery after photobleaching (FRAP)

Oocytes expressing Ddx3x-GFP were used to perform FRAP assays. FRAP acquisitions were performed on a Plan-APO ×60/1.4 NA objective on a spinning disk confocal microscope (Nikon Eclipse Ti, Nikon) equipped with a spinning-disk head (X1, Yokogawa), an Evolve EMCCD camera (Photometrics) and a thermostatic chamber (37°C, 5% CO2). The iLAS 2 module (Gataca System) was used to control the FRAP laser. Excitation was carried out on a fixed circular region of ∼10 µm of diameter and three equally sized regions in the surrounding cytoplasm. High intensity laser pulses (100% of 488 nm laser power, 20 pulses for a total of 0,48 ms of stimulation) were applied to photobleach fluorescence in these regions. Fluorescence recovery was then recorded in time-lapse mode with an interval of 500 ms over a total duration of 300 s. A pre-acquisition phase of 10 timepoints was used to establish a baseline fluorescence level. FRAP image sequences were realigned with Fiji StackReg plugin and regions of interest were selected for fluorescence recovery analysis. A spreadsheet was generated with the fluorescent intensity value of bleached regions in the cluster or cytoplasm at every timepoint, as well as of the whole cell and background. The spreadsheet was then uploaded on EasyFRAP-web (*51*). Recovery was measured as fluorescence intensity of photobleached regions corrected for background and normalized by the region’s prebleach and post-bleach intensities. Data was plotted on GraphPad 7 to obtain graph in Fig. 3E.

### Electron Microscopy

For each experimental condition, groups of about 10 oocytes per stage were adhered on a round glass coverslip (12 mm diameter, 150 µm thickness) previously coated with gelatin and polylysine with etched landmarks. Coverslips were prepared in a 24-well plate, at room temperature (except when mentioned). Samples were fixed 1 h with 2 % glutaraldehyde and 0.04 % Ruthenium Red in 0.1 M sodium cacodylate buffer (pH 7.4), washed with the same buffer, postfixed 1 h with 1 % osmium tetroxide and 1.5 % potassium ferrocyanide in 0.1 M cacodylate buffer at 4°C, and washed with ultrapure water. Samples were dehydrated at 4°C with ethanol (50-70-95-3x100 %, 5 to 15 min each step) and infiltrated during 24 hours with resin (25-50-2x100 %, Agar 100 kit, last step in vacuum). Coverslips were adhered on glass slides with a drop of resin and covered with upside-down gelatin capsules filled with resin, and were polymerized 48 h at 60°C. Capsules were then detached with the heat of a lighter under the glass slides (*52*). Blocks were cut with an ultramicrotome (Ultracut UCT, Leica microsystems) and a diamond knife (Ultra 35°, Diatome) until reaching the plane of maximum diameter of the oocyte. 80 nm thin sections were deposited on 5 mm square silicon wafers and were contrasted 15 min with 2.5 % uranyl acetate. Wafers were stuck on aluminum stubs with silver paint. Sections were observed with a Field Emission Scanning Electron Microscope (Gemini 500, Zeiss), operating in high vacuum, at 1.5 kV, with the high current mode and a 60 μm aperture diameter (corresponding to a specimen current of 411 pA), and a working distance around 2 mm. An in-column detector was used to collect backscattered electrons (filtering grid at 600 V). LookUp Table was inverted to obtain transmission electron microscope-like images. Automated acquisitions were performed using Atlas 5 (Fibics), with a pixel dwell time of 51.2 μs, a pixel size of 3 nm, an image definition of 7k x 7k pixels, and an overlap of 15 % between images. Full maps of oocytes (6 oocytes per stage) were obtained after stitching the image mosaics.

### RNA sequencing and differential gene expression analysis

A total of 10 C57BL/6J female mice (Charles River laboratories) aged four weeks were used for the experiments. Oocytes at the Cluster and Intermediary stage were separated in 2 groups of 3 x 30 oocytes per sample and collected in 2 µl PBS; 3 biological replicates were analyzed for each condition. Total RNA was extracted using the RNAqueous Micro Total RNA Isolation Kit (Thermofisher, AM1931). Oocytes were washed 3 times in PBS, resuspended in lysis buffer supplemented with 1% of ß-Mercaptoethanol, frozen in liquid nitrogen and conserved at -80°C overnight. After unfreezing, RNA extraction was carried out following the manufacturer’s instructions. RNA was eluted twice in 10 μl elution buffer. Samples were then treated with DNAse I and stored at -80°C. For cDNA libraries preparation, 300 ng of total RNA were amplified and converted to cDNA using SMART-Seq v4 Ultra Low Input RNA kit (Clontech) with 16 PCR cycles. Afterwards an average of 400 pg of amplified cDNA was used to prepare library following Nextera XT DNA kit (Illumina). Libraries were multiplexed by 6 on a P3 flow cell. A 68 bp read sequencing was performed on a NextSeq 2000 device (Illumina). A mean of 210 ± 28 million passing Illumina quality filter reads was obtained for each of the 6 samples. The analyses were performed using the Eoulsan pipeline (*53*), including read filtering, mapping, alignment filtering, read quantification, normalization and differential analysis. Before mapping, poly N read tails were trimmed, reads ≤40 bases were removed, and reads with quality mean ≤30 were discarded. Reads were then aligned against the *Mus musculus* genome from Ensembl version 105 using STAR (version 2.7.8a) (*54*). Alignments from reads matching more than once on the reference genome were removed using Java version of samtools (*55*). To compute gene expression, Mus musculus GFF3 and GTF genome annotation from Ensembl version 105 database was used. All overlapping regions between alignments and referenced exons (or genes) were counted and aggregated by genes using HTSeq-count 0.5.3 (*56*). The sample counts were normalized using DESeq2 1.8.1 (*57*). Statistical treatments and differential analyses were also performed using DESeq2 1.8.1. The RNASeq gene expression data and raw fastq files are available on the EMBL-EBI repository (https://www.ebi.ac.uk/) under ArrayExpress accession number: E-MTAB- 14892.

### Gene enrichment analyses

Data obtained from mass spectrometry were analysed with ShinyGO V0.81 (*58*) on the total list of proteins identified in the isolated cluster and using the list of proteins identified in the whole Cluster stage oocyte as background. To identify possible prion-like domains, primary sequences of proteins identified by mass spectrometry were analysed using PLAAC (Prion-Like Amino Acid Composition, (*50*)).The same software was used to generate protein sequence representation in Fig. S2A.

For the GSEA analysis, raw pooled count matrices were used from RNAseq data. The ’DESeq2’ R library (*57*) was used to import raw counts into a ’DESeqDataSet’. Empty rows were removed and differential expression analysis was conducted with the ’DESeq’ function. Fold changes were shrunken with the ’lfcShrink’ function, before the results were extracted and ordered by descending log2 fold change. The ordered list was then passed on to ’clusterProfiler’ R library to conduct GSEA Gene Ontology pathway analysis (*59*). The GSEA (*60*) was conducted for the same ordered list at the molecular functions (MF) level (Fig. 4B and 4D) and biological processes (BP) level (Fig. 4C). GSEA analysis was run with the ’fgsea’ underlying R method (*61*). Genes were annotated for the ENSEMBL annotation using the specific mus musculus ’org.Mm.eg.db’ R library. Dot plots were generated respectively with the top 8 and 10 most enriched pathways. Pathways were ordered by gene ratio, i.e. the percentage of genes, in each adequate Gene Ontology pathway, recognized as enriched by the GSEA analysis. P-values were adjusted by the Benjamini-Hochberg procedure (FDR). “Count” indicates the number of genes in each pathway on the y axis.

### Nascent protein synthesis using the OPP kit

The Click-iT^®^ Plus OPP Alexa Fluor^®^ 647 Protein Synthesis Assay Kit (Thermofisher, C10458) was used for the detection of protein synthesis. Prophase I oocytes at the Cluster stage were treated or not with 100 μg/ml puromycin for 1 hour before incubation with the puromycin analog OPP (O-propargyl-puromycin) 10 μM for 30 minutes and prior to their fixation. OPP detection was carried out following manufacturer’s instructions.

### Oocytes segmentation and feature extraction using Oocytor

The pipeline based on Oocytor (*62*) was applied to transmitted light images acquired for dry-mass measurement. As a first step, Oocytor detects the oocyte cortex and zona pellucida contours from transmitted light images based on neural networks (with a U-Net architecture, (*63*)). A model previously trained on images from both mouse and human oocyte was used (accessible here: https://github.com/gletort/Oocytor/tree/main/models/cortex/full for cortex segmentation and https://github.com/gletort/Oocytor/tree/main/models/zp for zona pellucida segmentation). Dry-mass raw images contain the refractive index information in multiple angles and present a checkboard-like pattern at the pixels level. At the full image level, the main components of the oocyte are nevertheless visible so we were able to run Oocytor directly on these images without having to retrain the networks on these specific types of images. However, as the neural networks were not fine-tuned, all the results were manually checked and about one third were badly segmented. We preprocessed the imaged for which the segmentation failed by readjusting the crop, converting the image to 8-bit, smoothing the pixel values (“Smooth” function of Fiji), shifting the pixel value distribution to a mean of 100 and sometimes with contrast enhancing (“Enhance Contrast” function of Fiji). These preprocessing steps allowed to make the images more similar to the training dataset. Oocytor was run again on the reprocessed images and gave better segmentation results. In a second step we extracted morphological values describing the oocyte. Oocytor proposes features either based on the segmentation contour or on the intensity distribution. Here as the pixel values reflect different refractive indexes and the images were not all preprocessed in the same way, we did not analyze the intensity-based measures.

### Dry mass measurement

A commercial PHASICS SID4-sC8 wave front sensor, recording phase images with up to 720x720 pixels (pixel size of 19.5 µm) was used. The dry mass of oocytes was measured according to (*64*). To image the oocytes, the aperture diaphragm was closed to the smallest size, in order to ensure that the sample was illuminated by a highly coherent and plane wave. A reference wave front was acquired close to the oocyte to measure. The Oocytor plugin (*62*) was used to segment oocytes from the interferograms acquired with the wave front sensor. A 4^th^-order polynomial fit of the optical path difference (OPD) values was made and subtracted to the whole phase images, including the oocyte pixels. The resulting OPD values inside the oocyte ROI determined by the Oocytor plugin were summed to extract the dry mass. The oocyte mean diameter was deduced from the ROI area, assuming the oocyte projection in the equatorial plane as a disk.

### Image analysis and quantifications

All images were analysed on Fiji (Version 2.1.0/1.53c, (*65*)).

Quantification of oocytes and Zollo body surface in Fig. 1B was obtained from live oocytes injected or not with Ddx3x-GFP construct. For the quantification of fluorescence intensity ratio between Zollo body and cytoplasm in Fig. 3C, 5C, 5D, S2D, the ROIs outlining the Zollo body, the nucleus and the whole oocyte were defined for each oocyte based on the signal from Mito Tracker, Tomm20 or autofluorescence of organelles at 488 nm. Cytoplasmic signal was obtained on the whole oocyte ROI excluding signal from the Zollo body and the nucleus, using the “XOR” function in Fiji. Signal intensity was measured after background removal on Fiji. Ratio of the signal intensity in the Zollo body vs cytoplasm was calculated on Excel and plotted on GraphPad 7. Phospho-mTOR intensity in Fig. 5F corresponds to the mean grey value of the maximal intensity z projection for the whole oocyte. Quantifications of zona pellucida width in Fig. S1B were obtained with Oocytor (*62*). Quantification of the number of oocytes with or without the cluster at different mouse ages from Fig. S1C was performed on live oocytes stained with Mito Tracker and/or SYTO RNA Select. Quantification of Zollo bodies dissolved by Nocodazole treatment in Fig. S1E was estimated on data from assays as in Fig. 5E-F (for 2 h) and S1G-H (for 30 min). Quantification of Zb size before and after Nocodazole treatment from Fig. S1F was performed on oocytes treated for 2 hours as in Fig. 5E-F.

### Statistical analysis

All graphs and statistical analyses were generated using MS Excel (Version 16.78) and GraphPad Prism 7 or 9. SD was used as y-axis error bars for bar charts and curves plotted from the mean value of the data. Statistical significance based on one way ANOVA followed by Tukey’s post hoc test, unpaired t test with Welch’s correction or Mann-Whitney test was calculated in Graphpad Prism 7. Linear regression analysis was performed in GraphPad Prism 9. All data are from at least three independent experiments. In the figures, significance is designated as follows: ****= p < 0.0001, ***= p < 0.001, **p= < 0.01, *= p < 0.05. Nonsignificant values are indicated as n.s.

---

**Fig. S1.**
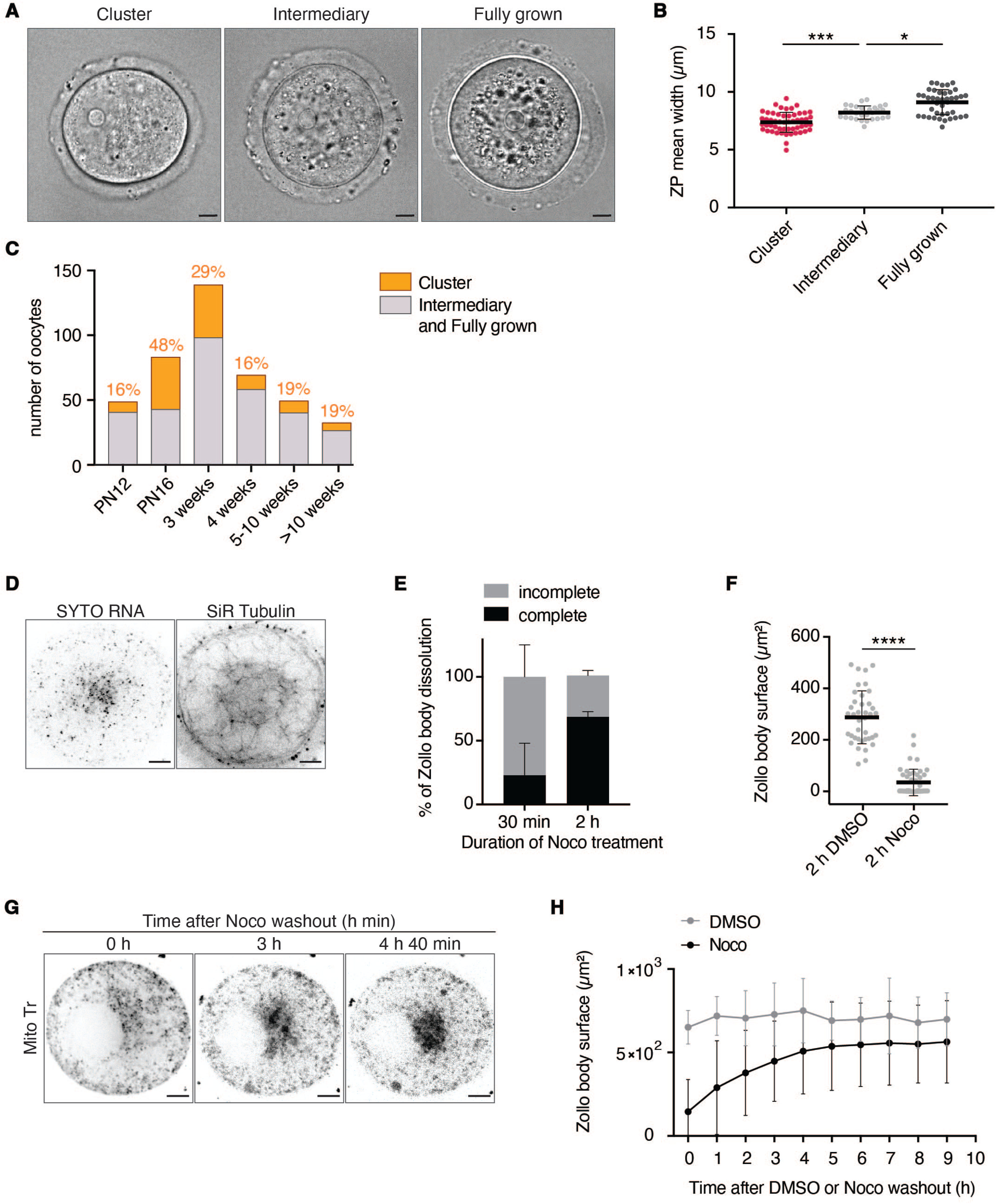
The Zb is found at a specific stage in oocyte growth and its integrity is linked to microtubules. **(A)** Transmitted light images of oocyte at the Cluster (left), Intermediary (middle) and Fully grown (right) stages. **(B)** Quantification of Zona Pellucida (ZP) thickness for oocytes at the Cluster, Intermediary or Fully grown stage, performed with the AI tool Oocytor. N=53 Cluster, N=29 Intermediary, N=41 Fully grown. **(C)** Proportion of oocytes at Cluster vs Intermediary or Fully grown stages at different mouse ages. N=78 oocytes, 2 mice for PN12; N=333 oocytes, 4 mice for PN16; N=555 oocytes, 4 mice for 3 weeks; N=415 oocytes, 6 mice for 4 weeks; N=197 oocytes, 4 mice for 5-10 weeks; 162 oocytes, 5 mice for >10 weeks. **(D)** Representative images of a Cluster stage oocyte stained with SYTO RNA (left) and SiR Tubulin (right). **(E)** Percentage of oocytes at the Cluster stage whose Zb dissolved completely or not after 30 min or 2 h treatment with 1 μM Nocodazole (Noco); N= 70 for 30 min treatment, 6 separate experiments; N=45 for 2 h, 5 separate experiments. **(F)** Quantification of Zb surface in oocytes at the Cluster stage treated with 1 μM Noco or its control DMSO for 2 h from (E); N=39 for DMSO, N=45 for Noco. (**G)** Representative images of a Cluster stage oocyte stained with Mito Tr, treated 30 min with 1 μM Nocodazole, then washed out and observed at different time points. **(H)** Quantification of Zb reformation after Noco washout as in (G). Only oocytes with completely dissolved Zb were selected for quantification after Noco washout. N=15 oocytes, 4 mice for Noco treatment; N=9 oocytes, 3 mice for DMSO control. Data are shown as mean ± SD. P values were calculated using one-way ANOVA with Tukey’s post hoc test. Scale bars 10 μm. ns = not significant; * = p<0.05; ** = p<0.01; *** = p<0.001; **** = p<0.0001.

**Fig. S2.**
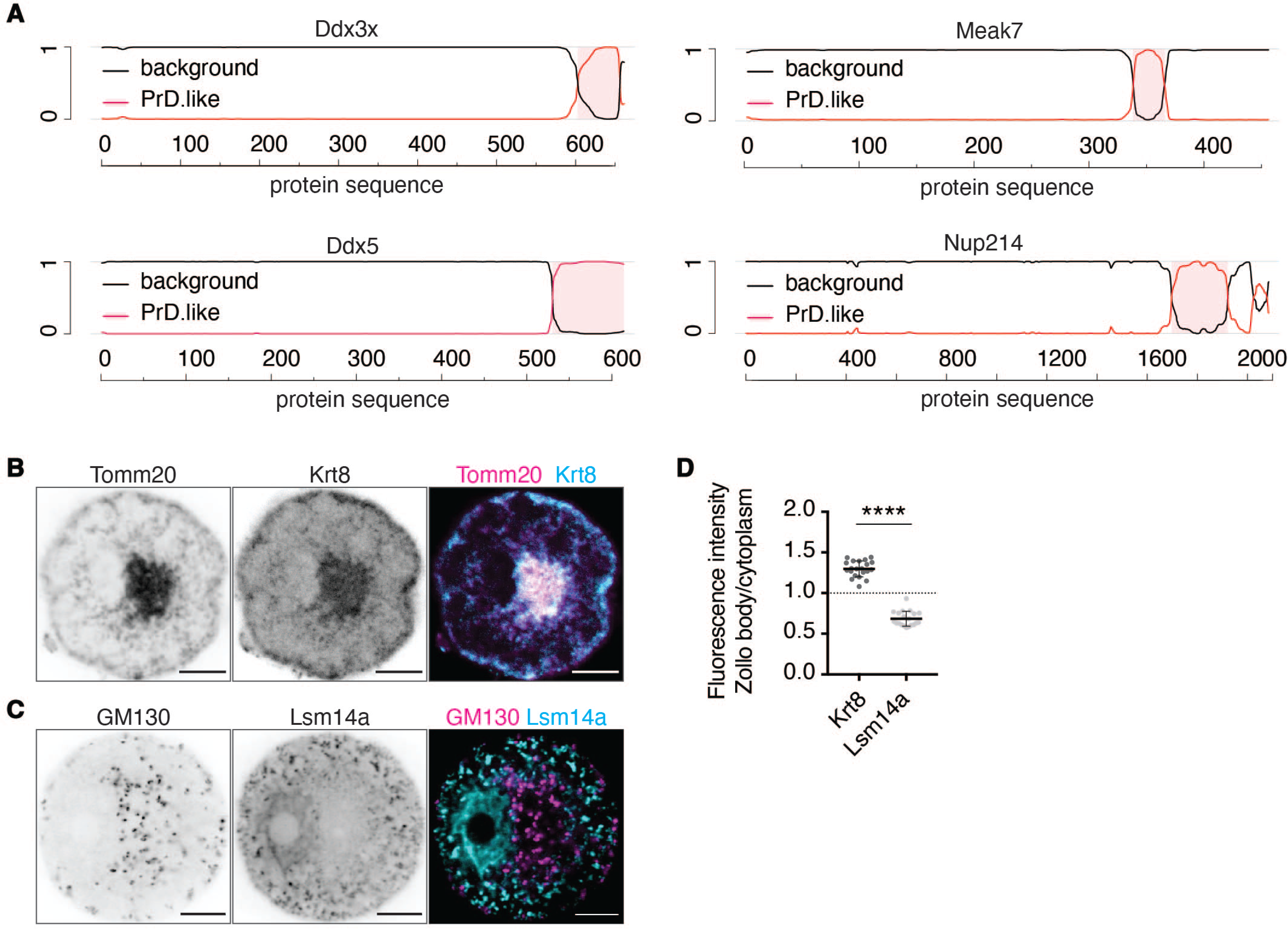
The Zb contains prion-like proteins but not the canonical RNP marker Lsm14a. **(A)** PLAAC algorithm results predicting prion-like domains in proteins enriched in the Zb. Pink shading indicates predicted prion-like sequences. **(B)** Staining indicating enrichment of Keratin 8 (Krt8, middle) in the Zb, identified by mitochondria (Tomm20, left). The right panel shows the merge of mitochondria (pink) and Krt8 (blue) staining. **(C)** Staining indicating the absence of Lsm14a staining (middle) inside the Zb, identified by Golgi (GM130, left). The right panel shows the merge of mitochondria (pink) and Lsm14a (blue) staining. **(D)** Quantification of Krt8 and Lsm14a signal intensities in the Zb compared to the cytoplasm from (B) and (C). N=21 for Krt8, 3 separate experiments; N=20 for Lsm14a, 3 separate experiments. Data are shown as mean ± SD. P values were calculated using unpaired two-tailed Student’s t test. Scale bars 10 μm. ns = not significant; * = p<0.05; ** = p<0.01; *** = p<0.001; **** = p<0.0001.

**Fig. S3.**
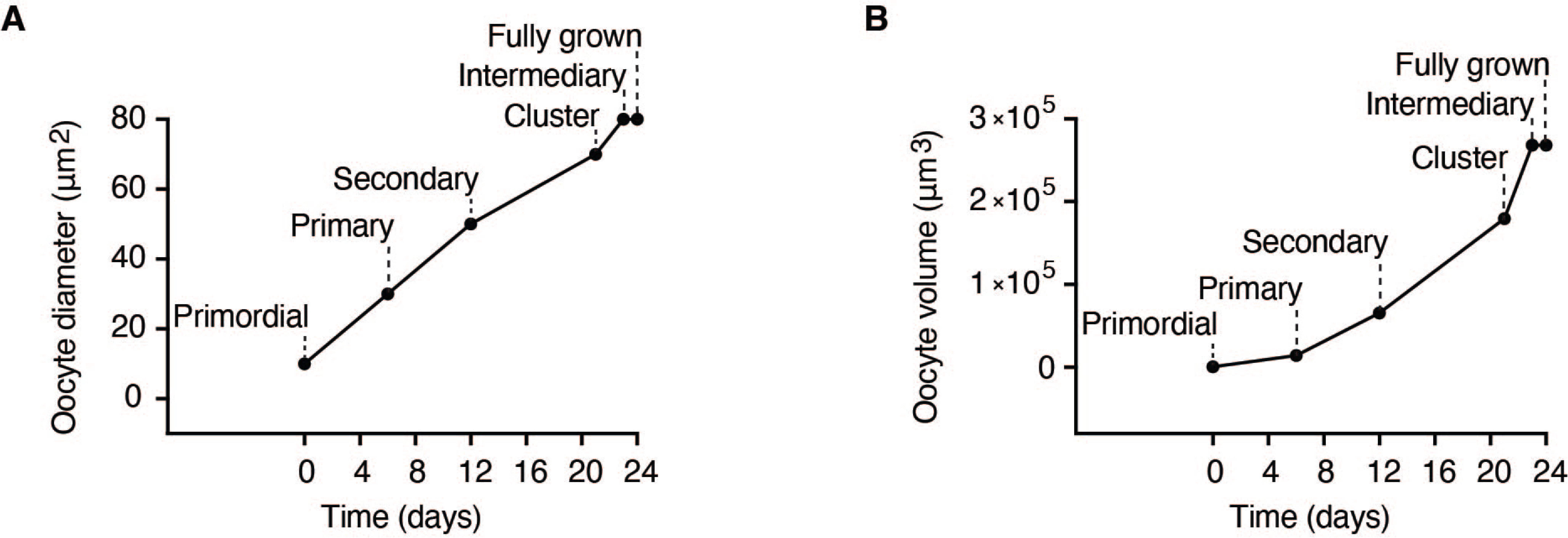
Analysis of previous literature reveals that the Cluster stage marks a phase in which oocyte growth is accelerated. **(A)** Schematic representation of the information found in (*3*, *27–31*), regarding oocytes diameter at the Primordial, Primary, Secondary, Cluster, Intermediary and Fully grown stage. **(B)** Schematic representation of the information found in (*3*, *27–31*), regarding oocytes volume at the Primordial, Primary, Secondary, Cluster, Intermediary and Fully grown stage.

## References

1. C. Bouniol-Baly, L. Hamraoui, J. Guibert, N. Beaujean, M. S. Szöllösi, P. Debey, Differential transcriptional activity associated with chromatin configuration in fully grown mouse germinal vesicle oocytes. Biol. Reprod. 60, 580–587 (1999).

2. M. Almonacid, A. Al Jord, S. El-Hayek, A. Othmani, F. Coulpier, S. Lemoine, K. Miyamoto, R. Grosse, C. Klein, T. Piolot, P. Mailly, R. Voituriez, A. Genovesio, M.-H. Verlhac, Active Fluctuations of the Nuclear Envelope Shape the Transcriptional Dynamics in Oocytes. Dev. Cell 51, 145–157.e10 (2019).

3. A. Al Jord, G. Letort, S. Chanet, F.-C. Tsai, C. Antoniewski, A. Eichmuller, C. Da Silva, J.-R. Huynh, N. S. Gov, R. Voituriez, M.-É. Terret, M.-H. Verlhac, Cytoplasmic forces functionally reorganize nuclear condensates in oocytes. Nat. Commun. 13, 5070 (2022).

4. M. Kloc, S. Bilinski, L. D. Etkin, The Balbiani body and germ cell determinants: 150 years later. Curr. Top. Dev. Biol. 59, 1–36 (2004).

5. A. Jamieson-Lucy, M. C. Mullins, The vertebrate Balbiani body, germ plasm, and oocyte polarity. Curr. Top. Dev. Biol. 135, 1–34 (2019).

6. E. Boke, M. Ruer, M. Wühr, M. Coughlin, R. Lemaitre, S. P. Gygi, S. Alberti, D. Drechsel, A. A. Hyman, T. J. Mitchison, Amyloid-like Self-Assembly of a Cellular Compartment. Cell 166, 637–650 (2016).

7. A. H. Jamieson-Lucy, M. Kobayashi, Y. James Aykit, Y. M. Elkouby, M. Escobar- Aguirre, C. E. Vejnar, A. J. Giraldez, M. C. Mullins, A proteomics approach identifies novel resident zebrafish Balbiani body proteins Cirbpa and Cirbpb. Dev. Biol. 484, 1–11 (2022).

8. F. L. Marlow, M. C. Mullins, Bucky ball functions in Balbiani body assembly and animal-vegetal polarity in the oocyte and follicle cell layer in zebrafish. Dev. Biol. 321, 40–50 (2008).

9. H. Hwang, S. Chen, M. Ma, null Divyanshi, H.-C. Fan, E. Borwick, E. Böke, W. Mei, J. Yang, Solubility phase transition of maternal RNAs during vertebrate oocyte-to- embryo transition. Dev. Cell 58, 2776–2788.e5 (2023).

10. S. Cheng, G. Altmeppen, C. So, L. M. Welp, S. Penir, T. Ruhwedel, K. Menelaou, K. Harasimov, A. Stützer, M. Blayney, K. Elder, W. Möbius, H. Urlaub, M. Schuh, Mammalian oocytes store mRNAs in a mitochondria-associated membraneless compartment. Science 378, eabq4835 (2022).

11. M. E. Pepling, J. E. Wilhelm, A. L. O’Hara, G. W. Gephardt, A. C. Spradling, Mouse oocytes within germ cell cysts and primordial follicles contain a Balbiani body. Proc. Natl. Acad. Sci. U.S.A. 104, 187–192 (2007).

12. L. Dhandapani, M. C. Salzer, J. M. Duran, G. Zaffagnini, C. De Guirior, M. A. Martínez-Zamora, E. Böke, Comparative analysis of vertebrates reveals that mouse primordial oocytes do not contain a Balbiani body. J. Cell. Sci. 135, jcs259394 (2022).

13. G. E. Neurohr, R. L. Terry, J. Lengefeld, M. Bonney, G. P. Brittingham, F. Moretto, T. P. Miettinen, L. P. Vaites, L. M. Soares, J. A. Paulo, J. W. Harper, S. Buratowski, S. Manalis, F. J. van Werven, L. J. Holt, A. Amon, Excessive Cell Growth Causes Cytoplasm Dilution And Contributes to Senescence. Cell 176, 1083–1097.e18 (2019).

14. S. Manohar, M. E. Estrada, F. Uliana, K. Vuina, P. M. Alvarez, R. A. M. de Bruin, G. E. Neurohr, Genome homeostasis defects drive enlarged cells into senescence. Mol. Cell 83, 4032–4046.e6 (2023).

15. I.-W. Lee, A. P. Tazehkand, Z.-Y. Sha, D. Adhikari, J. Carroll, An aggregated mitochondrial distribution in preimplantation embryos disrupts nuclear morphology, function, and developmental potential. Proc. Natl. Acad. Sci. U. S. A. 121, e2317316121 (2024).

16. M. C. Pamula, R. Lehmann, How germ granules promote germ cell fate. Nat. Rev. Genet. 25, 803–821 (2024).

17. S. Kar, R. Deis, A. Ahmad, Y. Bogoch, A. Dominitz, G. Shvaizer, E. Sasson, A. Mytlis, A. Ben-Zvi, Y. M. Elkouby, The Balbiani body is formed by microtubule-controlled molecular condensation of Buc in early oogenesis. Curr. Biol. 35, 315–332.e7 (2025).

18. J. Azoury, K. W. Lee, V. Georget, P. Rassinier, B. Leader, M.-H. Verlhac, Spindle positioning in mouse oocytes relies on a dynamic meshwork of actin filaments. Curr. Biol. 18, 1514–1519 (2008).

19. M. Schuh, J. Ellenberg, A new model for asymmetric spindle positioning in mouse oocytes. Curr. Biol. 18, 1986–1992 (2008).

20. M. Luksza, I. Queguigner, M.-H. Verlhac, S. Brunet, Rebuilding MTOCs upon centriole loss during mouse oogenesis. Dev. Biol. 382, 48—56 (2013).

21. H. Zhang, T. Zhang, X. Wan, C. Chen, S. Wang, D. Qin, L. Li, L. Yu, X. Wu, LSM14B coordinates protein component expression in the P-body and controls oocyte maturation. J. Genet. Genom. 51, 48–60 (2024).

22. A. Hubstenberger, M. Courel, M. Bénard, S. Souquere, M. Ernoult-Lange, R. Chouaib, Z. Yi, J.-B. Morlot, A. Munier, M. Fradet, M. Daunesse, E. Bertrand, G. Pierron, J. Mozziconacci, M. Kress, D. Weil, P-Body Purification Reveals the Condensation of Repressed mRNA Regulons. Mol. Cell 68, 144–157.e5 (2017).

23. T. Hochstoeger, J. A. Chao, Towards a molecular understanding of the 5′TOP motif in regulating translation of ribosomal mRNAs. Semin. Cell Dev. Biol. 154, 99–104 (2024).

24. J. Liu, Y. Xu, D. Stoleru, A. Salic, Imaging protein synthesis in cells and tissues with an alkyne analog of puromycin. Proc. Natl. Acad. Sci. U.S.A. 109, 413–418 (2012).

25. G. Y. Liu, D. M. Sabatini, mTOR at the nexus of nutrition, growth, ageing and disease. Nat. Rev. Mol. Cell Biol. 21, 183–203 (2020).

26. R. Rollin, J.-F. Joanny, P. Sens, Physical basis of the cell size scaling laws. eLife 12, e82490 (2023).

27. J. J. Eppig, K. Wigglesworth, F. L. Pendola, The mammalian oocyte orchestrates the rate of ovarian follicular development. Proc. Natl. Acad. Sci. U. S. A. 99, 2890–2894 (2002).

28. H. Clarke, Control of Mammalian Oocyte Development by Interactions with the Maternal Follicular Environment. Results Probl. Cell. Differ. 63, 17–41 (2017).

29. J. Griffin, B. R. Emery, I. Huang, C. M. Peterson, D. T. Carrell, Comparative analysis of follicle morphology and oocyte diameter in four mammalian species (mouse, hamster, pig, and human). J. Exp. Clin. Assist. Reprod. 3, 2 (2006).

30. J. J. Eppig, M. J. O’Brien, Development in vitro of mouse oocytes from primordial follicles. Biol. Reprod. 54, 197–207 (1996).

31. 31. S. F. Gilbert, “Oogenesis” in Developmental Biology. 6th Edition (Sinauer Associates, 2000).

32. R. Chen, W. Stainier, J. Dufourt, M. Lagha, R. Lehmann, Direct observation of translational activation by a ribonucleoprotein granule. Nat. Cell Biol. 26, 1322–1335 (2024).

33. M.-C. Lai, Y.-H. W. Lee, W.-Y. Tarn, The DEAD-box RNA helicase DDX3 associates with export messenger ribonucleoproteins as well as tip-associated protein and participates in translational control. Mol. Biol. Cell 19, 3847–3858 (2008).

34. E. Nikalayevich, G. Letort, G. de Labbey, E. Todisco, A. Shihabi, H. Turlier, R. Voituriez, M. Yahiatene, X. Pollet-Villard, M. Innocenti, M. Schuh, M.-E. Terret, M.-H. Verlhac, Aberrant cortex contractions impact mammalian oocyte quality. Dev. Cell, S1534-5807(24)00047–9 (2024).

35. A. Hubstenberger, S. L. Noble, C. Cameron, T. C. Evans, Translation Repressors, an RNA Helicase, and Developmental Cues Control RNP Phase Transitions during Early Development. Dev. Cell 27, 161–173 (2013).

36. E. Boke, T. J. Mitchison, The balbiani body and the concept of physiological amyloids. Cell Cycle 16, 153–154 (2017).

37. F. Buttgereit, M. D. Brand, A hierarchy of ATP-consuming processes in mammalian cells. Biochem. J. 312 **(Pt** **1**), 163–167 (1995).

38. F. Bontems, A. Stein, F. Marlow, J. Lyautey, T. Gupta, M. C. Mullins, R. Dosch, Bucky ball organizes germ plasm assembly in zebrafish. Curr. Biol. 19, 414–422 (2009).

39. J.-W. Shih, W.-T. Wang, T.-Y. Tsai, C.-Y. Kuo, H.-K. Li, Y.-H. Wu Lee, Critical roles of RNA helicase DDX3 and its interactions with eIF4E/PABP1 in stress granule assembly and stress response. Biochem. J. 441, 119–129 (2012).

40. B. Shi, J. Heng, J.-Y. Zhou, Y. Yang, W.-Y. Zhang, M. J. Koziol, Y.-L. Zhao, P. Li, F. Liu, Y.-G. Yang, Phase separation of Ddx3xb helicase regulates maternal-to-zygotic transition in zebrafish. Cell Res. 32, 715–728 (2022).

41. M. C. Owens, H. Shen, A. Yanas, M. S. Mendoza-Figueroa, E. Lavorando, X. Wei, H. Shweta, H.-Y. Tang, Y. E. Goldman, K. F. Liu, Specific catalytically impaired DDX3X mutants form sexually dimorphic hollow condensates. Nat. Commun. 15, 9553 (2024).

42. M. de Castro Fonseca, J. F. de Oliveira, B. H. S. Araujo, C. Canateli, P. F. V. do Prado, D. P. Amorim Neto, B. P. Bosque, P. V. Rodrigues, J. V. P. de Godoy, K. Tostes, H. V. R. Filho, A. F. Z. Nascimento, A. Saito, C. C. C. Tonoli, F. A. H. Batista, P. S. L. de Oliveira, A. C. Figueira, S. Souza da Costa, A. C. V. Krepischi, C. Rosenberg, H. Westfahl, A. J. R. da Silva, K. G. Franchini, Molecular and cellular basis of hyperassembly and protein aggregation driven by a rare pathogenic mutation in DDX3X. iScience 24, 102841 (2021).

43. B. Leader, H. Lim, M. J. Carabatsos, A. Harrington, J. Ecsedy, D. Pellman, R. Maas, P. Leder, Formin-2, polyploidy, hypofertility and positioning of the meiotic spindle in mouse oocytes. Nat. Cell Biol. 4, 921–928 (2002).

44. M. Verlhac, J. Kubiak, H. Clarke, B. Maro, Microtubule and chromatin behavior follow MAP kinase activity but not MPF activity during meiosis in mouse oocytes. Development 120, 1017—1025 (1994).

45. A. Reis, H.-Y. Chang, M. Levasseur, K. T. Jones, APCcdh1 activity in mouse oocytes prevents entry into the first meiotic division. Nat. Cell Biol. 8, 539–540 (2006).

46. M. Manil-Ségalen, M. Łuksza, J. Kanaan, V. Marthiens, S. I. R. Lane, K. T. Jones, M.-E. Terret, R. Basto, M.-H. Verlhac, Chromosome structural anomalies due to aberrant spindle forces exerted at gene editing sites in meiosis. J. Cell Biol. 217, 3416–3430 (2018).

47. M. Breuer, A. Kolano, M. Kwon, C.-C. Li, T.-F. Tsai, D. Pellman, S. Brunet, M.-H. Verlhac, HURP permits MTOC sorting for robust meiotic spindle bipolarity, similar to extra centrosome clustering in cancer cells. J. Cell Biol. 191, 1251—1260 (2010).

48. M.-H. Verlhac, C. Lefebvre, P. Guillaud, P. Rassinier, B. Maro, Asymmetric division in mouse oocytes: with or without Mos. Curr. Biol. 10, 1303–1306 (2000).

49. G. Zaffagnini, S. Cheng, M. C. Salzer, B. Pernaute, J. M. Duran, M. Irimia, M. Schuh, E. Böke, Mouse oocytes sequester aggregated proteins in degradative super- organelles. Cell 187, 1109–1126.e21 (2024).

50. A. K. Lancaster, A. Nutter-Upham, S. Lindquist, O. D. King, PLAAC: a web and command-line application to identify proteins with prion-like amino acid composition. Bioinformatics 30, 2501–2502 (2014).

51. G. Koulouras, A. Panagopoulos, M. A. Rapsomaniki, N. N. Giakoumakis, S. Taraviras, Z. Lygerou, EasyFRAP-web: a web-based tool for the analysis of fluorescence recovery after photobleaching data. Nucleic Acids Res. 46, W467–W472 (2018).

52. I. Hurbain, M. Romao, P. Bergam, X. Heiligenstein, G. Raposo, Analyzing Lysosome-Related Organelles by Electron Microscopy. Methods Mol. Biol. 1594, 43–71 (2017).

53. L. Jourdren, M. Bernard, M.-A. Dillies, S. Le Crom, Eoulsan: a cloud computing- based framework facilitating high throughput sequencing analyses. Bioinformatics 28, 1542–1543 (2012).

54. A. Dobin, C. A. Davis, F. Schlesinger, J. Drenkow, C. Zaleski, S. Jha, P. Batut, M. Chaisson, T. R. Gingeras, STAR: ultrafast universal RNA-seq aligner. Bioinformatics 29, 15–21 (2013).

55. H. Li, B. Handsaker, A. Wysoker, T. Fennell, J. Ruan, N. Homer, G. Marth, G. Abecasis, R. Durbin, 1000 Genome Project Data Processing Subgroup, The Sequence Alignment/Map format and SAMtools. Bioinformatics 25, 2078–2079 (2009).

56. S. Anders, P. T. Pyl, W. Huber, HTSeq—a Python framework to work with high- throughput sequencing data. Bioinformatics 31, 166–169 (2015).

57. M. I. Love, W. Huber, S. Anders, Moderated estimation of fold change and dispersion for RNA-seq data with DESeq2. Genome Biol. 15, 550 (2014).

58. S. X. Ge, D. Jung, R. Yao, ShinyGO: a graphical gene-set enrichment tool for animals and plants. Bioinformatics 36, 2628–2629 (2020).

59. G. Yu, L.-G. Wang, Y. Han, Q.-Y. He, clusterProfiler: an R package for comparing biological themes among gene clusters. OMICS 16, 284–287 (2012).

60. A. Subramanian, P. Tamayo, V. K. Mootha, S. Mukherjee, B. L. Ebert, M. A. Gillette, A. Paulovich, S. L. Pomeroy, T. R. Golub, E. S. Lander, J. P. Mesirov, Gene set enrichment analysis: a knowledge-based approach for interpreting genome-wide expression profiles. Proc. Natl. Acad. Sci. U.S. A. 102, 15545–15550 (2005).

61. A. A. Sergushichev, An algorithm for fast preranked gene set enrichment analysis using cumulative statistic calculation. bioRxiv 060012 [Preprint] (2016). 10.1101/060012.

62. G. Letort, A. Eichmuller, C. Da Silva, E. Nikalayevich, F. Crozet, J. Salle, N. Minc, E. Labrune, J.-P. Wolf, M.-E. Terret, M.-H. Verlhac, An interpretable and versatile machine learning approach for oocyte phenotyping. J. Cell Sci. 135, jcs260281 (2022).

63. O. Ronneberger, P. Fischer, T. Brox, U-Net: Convolutional Networks for Biomedical Image Segmentation. arXiv arXiv:1505.04597 [Preprint] (2015). 10.48550/arXiv.1505.04597.

64. S. Aknoun, J. Savatier, P. Bon, F. Galland, L. Abdeladim, B. Wattellier, S. Monneret, Living cell dry mass measurement using quantitative phase imaging with quadriwave lateral shearing interferometry: an accuracy and sensitivity discussion. J. Biomed. Opt. 20, 126009 (2015).

65. 65. J. Schindelin, I. Arganda-Carreras, E. Frise, V. Kaynig, M. Longair, T. Pietzsch, S. Preibisch, C. Rueden, S. Saalfeld, B. Schmid, J.-Y. Tinevez, D. J. White, V. Hartenstein, K. Eliceiri, P. Tomancak, A. Cardona, Fiji: an open-source platform for biological-image analysis. Nat. Methods 9, 676–682 (2012).

